# Distinct stage-specific transcriptional states of B cells in human tonsillar tissue

**DOI:** 10.1101/2021.08.17.456535

**Authors:** Diego A. Espinoza, Carole Le Coz, Neil Romberg, Amit Bar-Or, Rui Li

**Affiliations:** Graduate Group in Immunology, Perelman School of Medicine, University of Pennsylvania, Philadelphia, PA, USA; Center for Neuroinflammation and Experimental Therapeutics, Perelman School of Medicine, University of Pennsylvania, Philadelphia, PA, USA; Department of Neurology, Perelman School of Medicine, University of Pennsylvania, Philadelphia, PA, USA; Division of Immunology and Allergy, Children’s Hospital of Philadelphia, Philadelphia, PA, USA; Department of Pediatrics, Perelman School of Medicine, Philadelphia, PA, USA

## Abstract

B cells within secondary lymphoid tissues encompass a diverse range of activation states and multiple maturation processes that reflect antigen recognition and transition through the germinal center (GC) reaction, in which mature B cells differentiate into memory and antibody-secreting cells (ASCs). Here, using single-cell RNA-seq, we identify distinct activation and maturation profiles of B cells within and outside the GC reaction in human secondary lymphoid tissue. In particular, we identify a distinct, previously uncharacterized *CCL4*/*CCL3* chemokine-expressing B-cell population with an expression pattern consistent with BCR/CD40 activation. Furthermore, we present a computational method leveraging regulatory network inference and pseudotemporal modeling to identify upstream transcription factor modulation along the GC to ASC maturation axis. Our dataset provides valuable insight into the diverse functional profiles and maturation processes that B cells undergo within secondary lymphoid tissues and will be a useful resource on which to base further studies into the B-cell immune compartment.

**Highlights:**

1. scRNA-seq of human tonsillar B cells identifies distinct activation and maturation phenotypes.
2. Identification of a chemokine-expressing B-cell population in the human tonsil with a BCR and CD40 co-stimulatory gene signature.
3. Transcription factor regulatory network analysis identifies MYC and REL as predicted regulators of chemokine expression in the chemokine-expressing B-cell population.
4. Trajectory inference with gene and regulatory network modeling implicates novel transcription factors in the GC-to-ASC transition.

## INTRODUCTION

Within secondary lymphoid tissues, mature B cells encounter antigen and, across multiple maturation processes incorporating the germinal center (GC) reaction, undergo a number of activation and fate-determining states as they ultimately differentiate into either antigen-specific memory B cells or antibody-secreting cells (ASCs). These intra-maturation states likely reflect a diversity of B-cell functions inclusive of, but likely also beyond, the well-characterized B-cell functions of humoral memory development and antibody production. Currently, the functional capabilities of B cells beyond antibody secretion remain incompletely understood, but are of interest given the growing recognition of antibody-independent roles of B cells in both normal immune responses and in autoimmune disease^1–4^. Previous insights into the secondary lymphoid tissue B-cell compartment and its dynamics have largely relied on a-priori defined phenotypic markers to identify and further characterize cells of interest. To this end, high-throughput phenotypic characterization of B cells through single-cell RNA-seq (scRNA-seq) provides an alternative venue by which the diverse phenotypic and functional states of B cells can be explored in a more unbiased manner within the lymphoid tissue. Such an approach would not only permit unbiased characterization of B-cell functional states but also help better define existing B-cell maturation processes at a higher, transcriptomic resolution. Advances in single-cell technology have already contributed to better understanding of B cell maturation, as demonstrated in a recent study that leveraged scRNA-seq along with repertoire analysis to better understand the interplay between antibody class-switching and gene expression in human tonsillar B cells^5^.

Here, we utilize scRNA-seq to better characterize the diversity of distinct B-cell activation and maturation states within the human tonsil. We identify a *CCL4/CCL3*-expressing B-cell population with an activation state consistent with BCR and CD40 co-stimulation and transcriptional processes likely associated with activity of the transcription factor MYC. Furthermore, we leverage regulatory network analyses to provide transcription-factor level insight into the differentiation of ASCs from GC B cells. This dataset provides valuable insight into the diverse functional profiles of B cells and their maturation processes and will be a useful resource on which to base further studies into the B-cell immune compartment in health and disease states.

## RESULTS

### scRNA-seq of human tonsillar B cells identifies distinct populations of germinal-center B cells, non-germinal-center B cells, and antibody-secreting cells

We sorted live CD3^-^/CD14^-^/CD19^+^ tonsillar B cells from human donors (n = 3, donor information in **Table S1**) and recovered single cell transcriptomes using the 10X Chromium 3’ scRNA-seq platform (Fig. 1a). Following data pre-processing (see ***Methods,*** Fig. S1a-c) to enrich for live B cells and to integrate data across three donors, we analyzed 45,376 B-cell transcriptomes. We performed marker gene identification using a one-vs-all approach for each cluster to identify ‘marker’ genes for each cluster (Fig. S2a, full marker list in **Table S2**). Based on prior work, we used the overall RNA-expression patterns of *CCR7*, *CD38*, and *MZB1* (Fig. 1b-c) to partition our clusters into 3 sets: (i) non-germinal-center B (non-GC) cell clusters (*CCR7*+/*CD38*-, defined by high levels of the tissue emigrant marker gene *CCR7*^6^ and low expression of the *CD38*^7^ human GC and plasma cell marker gene); (ii) germinal center B (GC) cell clusters (*CCR7*-/*CD38*+/*PRDM1*-, defined by high levels of CD38 and low levels of *PRDM1*); and (iii) antibody-secreting cell (ASC) clusters (*CCR7*-/*CD38*+/*PRDM1*+, defined by high levels of *CD38* and *PRDM1,* a plasma cell marker gene^8^, but low levels of *CCR7*).

**Figure 1:**
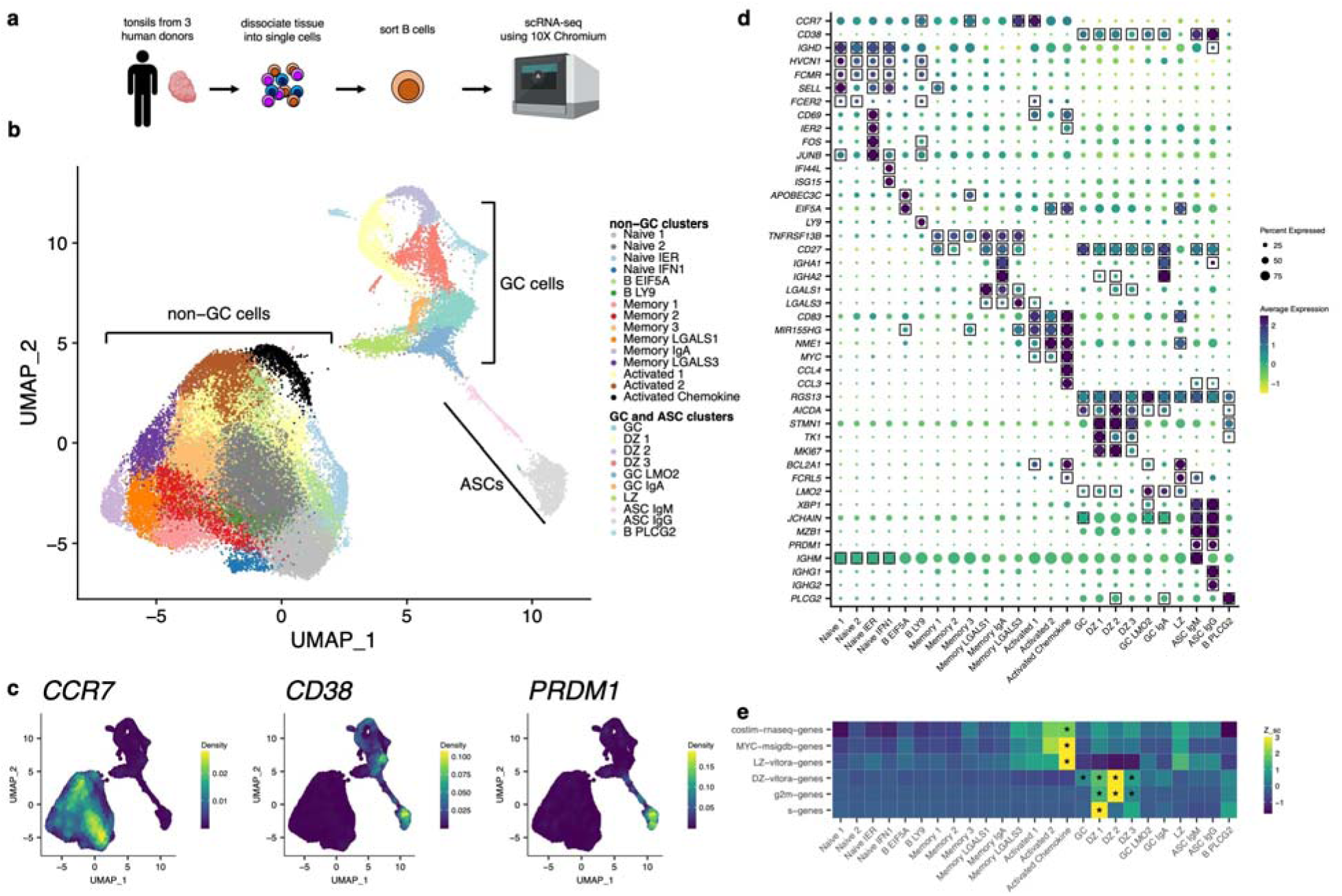
Single-cell RNA-sequencing of human tonsillar B cells identifies distinct B cell maturation and activation states. **(a)** Experimental design for tonsillar B-cell isolation, culture, and scRNA-seq. **(b)** UMAP projection of 45,376 single B-cell transcriptomes from 3 human donors. **(c)** Visualization of *CCR7*, *CD38*, and *MZB1* expression patterns in UMAP space using the kernel density estimation method in the *Nebulosa* R package. (**d**) Selected marker-gene average expression (scaled log normalized counts) for each cluster and proportion of cells with transcript detected. Genes were determined to be marker-genes if their average log2 fold-change was > 0.3 for the cluster of interest and adjusted p-value was < 0.01. Marker-gene expression is denoted by square boxes on gene-cluster pairs. **(e)** Selected scaled average AUCell scores for five genesets across each cluster. Heatmap cells were labelled with a star (*) if the geneset was enriched within that cluster (Wilcoxon rank-sum test, AUC > 0.85).

### Clusters of non-GC B cells capture multiple distinct states of B cell activation

We first focused on characterizing the transcriptional phenotypes of clusters within the non-GC B cell clusters. Using *IGHD, FCMR,* and *HVCN1*^9^ expression to define the naïve pool, we identified four B-cell subclusters (Naïve 1, Naïve 2, Naïve IER [immediate early response activated], and Naïve IFN1 [type-1 interferon activated], Fig. 1b). SELL, another naïve B-cell marker, was enriched at the gene expression level in the Naïve 1, Naïve IER, and Naïve IFN1 clusters (Fig. 1d). The Naïve IER cluster’s marker genes included activation markers *IER2, FOS, JUNB*, and *CD69* (Fig. 1d). CD69 is an early marker of B-cell activation, suggesting that cells within this cluster shared an expression pattern consistent with an early form of B cell activation. In contrast, the Naïve IFN1 cluster’s marker genes included interferon-stimulated genes such as *IFI44L, ISG15*, suggesting that cells within this cluster shared an expression pattern indicative of cellular type-1 interferon activation. A fifth cluster, B EIF5A, colocalized partially with the naïve clusters in the dataset (Fig. 1b); however, the B EIF5A cluster expressed no naïve-like or memory-like marker genes, and instead showed enrichment for genes involved in transcriptional regulation, such as *APOBEC3C* and *EIF5A* (Fig. 1d), suggesting a B-cell state with increased transcriptional activity. Proximal to the Naïve clusters in the UMAP space, we identified a cluster (B LY9) that shared marker genes with Naïve B cells (*HVCN1, FCMR,* and *FCER2*), but expressed uniquely high levels of *LY9* (Fig. 1d).

We next identified 6 memory B-cell clusters notable for enrichment of the marker gene *TNFRSF13B*^10^ (Fig. 1d). 5 of these 6 clusters (the exception being Memory 3) expressed CD27, another memory B-cell marker, as a marker gene as well. Limiting the differential expression analysis to these 6 memory B-cell clusters (Fig. S3), we found that the Memory 2 and Memory 3 showed relative enrichment for *IGHD* and *IGHM*, indicating that these may represent non-class-switched memory B cells. In contrast, Memory 1 showed enrichment for *IGHG1* and *IGHG3* (Fig. S2b), indicating that their heavy-chain expression was more representative of a class-switched memory identity. The Memory IgA cluster expressed marker genes *IGHA1* and *IGHA2*, identifying what is likely a mucosal B-cell memory cell subset (Fig. 1d). Two other memory B-cell clusters, Memory LGALS1 and Memory LGALS3, exhibited high expression of galectin transcripts *LGALS1* (encoding galectin-1) and *LGALS3* (encoding galectin-3), respectively. The B-cell specific roles of galectin-1 and galectin-3, particularly in memory B cells, have not been fully determined^11^.

Three clusters (Activated 1, Activated 2, and Activated Chemokine) expressed the same set of three cellular activation-associated genes as marker genes: *CD83*, *MIR155HG*, and *NFKB1* (Fig. 1b, d). CD83 surface expression is known to be upregulated on B cells in a number of activation contexts, including LPS, CD40, and IgM stimulation^12^. *MIR155HG* encodes the microRNA miR-155, which has also been shown to be upregulated on B cells upon TLR, CD40, and BCR stimulation^13^ in mice and upon BCR cross-linking in Ramos cells^14^ and shown to enhance BCR signaling in CD40-stimulated human B cells^15^. Additionally, miR-155 is downregulated in GC B cells by BCL6, a GC-specific transcription factor^16^, suggesting that cells within the Activated 1, Activated 2, and Activated Chemokine clusters are either cells not participating in or have yet to participate the germinal center reaction. Overall, the co-expression pattern of *CD83*, and *MIR155HG* in these clusters suggest that cells within these clusters may represent B cells receiving BCR and/or CD40 signaling prior to entry into the GC reaction. In contrast to the Activated 1 cluster, however, Activated 2 and Activated Chemokine expressed higher levels of *NME1* (Fig. 1b).

Next, in order to better determine the similarity of the three Activated clusters to B cells stimulated through the BCR alone (as a proxy for B cells encountering antigen but not receiving T cell help), CD40 alone (as a proxy for B cells encountering activated T cells expressing CD40L, but not encountering their antigen), or both the BCR and CD40 (as a proxy for B cells encountering both their antigen and receving costimulation through interaction with activated T cells), we stimulated peripheral-blood B cells in vitro (n = 3, donor information in **Table S3**) to derive gene signatures for each stimulus (BCR-only signature, CD40-only signature, co-stimulation signature, **Table S4**). We found that the Activated 2 and Activated Chemokine clusters showed enrichment for only one of our in-vitro stimulation signatures: the co-stimulation signature (Fig. 1e), corroborating our marker-gene observations. Of note, we also observed marker-gene expression of *MYC* in our Activated clusters (Fig. 1e); MYC is a transcription factor known to be necessary for GC formation^17,18^ and only expressed in a small subset of LZ cells in the GC^17,18^. Thus, given the expression of the activation markers above, the enrichment for a BCR/CD40 co-stimulation signature, and elevated MYC transcript expression, it is plausible that cells within these Activated clusters are representative of cells primed for entry into the GC reaction. The Activated Chemokine cluster, in addition to sharing marker gene expression of *CD83*, *MIR155HG*, *NFKB1*, and *MYC* and showing enrichment for our BCR/CD40 co-stimulation signature, also showed cluster-specific marker gene expression of the chemokine transcripts *CCL4* and *CCL3* (Fig. 1d). These chemokines are known to be secreted by B cells and are chemoattractant for T cells (in particular, regulatory T cells^3,4^). Enrichment analysis of literature-defined gene sets, including MYC-signature and light-zone (LZ) specific gene sets, found that the Activated Chemokine cluster had the highest enrichment for these gene sets, in line with its upregulation of activation-associated transcripts (Fig. 1e). The functional roles and necessity of this chemokine-expressing B-cell population has not yet been elucidated with relation to the GC reaction.

### Clusters of GC B cells capture multiple distinct states of B-cell maturation

We next characterized clusters corresponding to cells within the GC reaction. Cells were determined to be participating in the GC reaction if they were significantly enriched for the marker-genes *CD38* (save for one cluster) and *RGS13*, another marker shown to be enriched in germinal center B cells^19^ (Fig. 1d). We then characterized individual clusters representative of cellular subsets/states within the GC reaction. Using prior gene signatures for LZ and DZ genes^20^, we found three clusters, DZ 1, DZ 2, and DZ 3, with the highest enrichment for the DZ transcriptional program (Fig. 1e), which included genes such as *AICDA*, *STMN1*, *TK1*, and *MKI67* (Fig. 1d), These cells also represented actively cycling cells, a process that appeared to be accurately captured by a cyclical structure in the UMAP projection (Fig. 1d) with DZ 1 representing cells with gene expression most consistent with S-phase, DZ 2 representing cells with gene expression most consistent with G2/M-phase, and DZ 3 showing no particular skew towards S- or G2/M-phase gene expression (Fig. 1e). Clusters GC and GC IgA represented cells without clear enrichment for DZ or LZ programs and likely represent a mixture of cells in both programs (Fig. 1d-e) or an intermediate LZ-DZ state, with GC IgA showing enrichment for the *IGHA1* and *IGHA2* heavy chain genes. Cluster LZ showed relatively high enrichment for the LZ program (Fig. 1e) compared to other GC clusters. The LZ-markers *BCL2A1* and *FCRL5* were also enriched within this cluster (Fig. 1d). Notably, the Activated 2 and Activated Chemokine clusters also showed high enrichment for the LZ program and MYC gene set (Fig. 1e), while likely representing cells not in the GC reaction given their expression of *CCR7* and lack of *CD38* expression. The underlying transcriptional similarity between these two cellular phenotypes is likely because the LZ transcriptional program is expected to share intracellular signaling mechanisms with the overall B-cell activation transcriptional program, as some LZ B cells are expected to undergo BCR/CD40 co-stimulation. Finally, the GC LMO2 cluster showed enrichment for the germinal center DZ and LZ marker genes *AICDA* and *BCL2A1*, along with the gene *LMO2*, while expressing marker-genes *XBP1* and *JCHAIN* (albeit at a subtle, but statistically significant level), which are instead associated with antibody-secreting cells, suggesting an intermediate cellular state between GC and antibody-secreting phenotypes (Fig. 1d). Finally, another cluster, B PLCG2, showed particularly high marker-gene expression of the *PLCG2* gene and a number of histone genes (Fig. S2) while sharing expression programs with the DZ program and largely co-localizing with DZ cells in the UMAP projection (Fig. 1d). The high expression of histone transcripts, expected to be restricted to the nucleus, suggests that these cells likely represent dead or dying recently cell-cycling DZ cells whose cytoplasmic RNA has leaked to the extracellular environment resulting in enrichment of nucleus-associated transcripts.

### Clusters of antibody-secreting B cells are segregated by IgM or IgG heavy chain expression

We identified 2 clusters likely representing ASCs: ASC IgM and ASC IgG. These clusters both showed marker-gene expression of canonical ASC genes *XBP1*, *JCHAIN*, *MZB1*, and *PRDM1*, but differed in terms of their antibody isotype (Fig. 1d).

In summary, our scRNA-seq approach resolved multiple populations of B cells within the human tonsil at a transcriptional level, recapitulating heterogeneous naïve B cell, memory B cell, GC cell and ASC subsets, some with distinct signatures of activation and maturation. Of particular interest, we identified a cluster, Activated Chemokine, with high expression of *CCL4* and *CCL3* transcripts and a transcriptional profile consistent with BCR/CD40 activation, potentially representing a B-cell population secreting T-cell-attractant chemokines and interacting with T cells prior to entry into the GC within human tonsil.

### Transcription factor regulatory network analysis identifies MYC, REL, and FOSL1 as transcription factors driving the production of chemokine transcripts in the Activated Chemokine cluster

We next sought to use gene regulatory network (GRN) analysis to identify transcription factor signatures associated with each cluster in order to hypothesize the upstream transcriptional drivers of the observed cellular identities. Using the *SCENIC*^21^ pipeline, and a given list of curated transcription factors (TFs) and corresponding motif information, we derived transcription factor GRNs (*regulons*) from our scRNA-seq data. Heretofore, a *regulon* refers to a TF and its set of SCENIC-predicted downstream targets. We next inferred the single-cell *activity* of each regulon using the *AUCell* algorithm, which quantifies a measure of enrichment for a regulon’s predicted targets within the expressed genes for each cell. Differential activity of regulons was then identified between clusters by using a Wilcoxon rank-sum test (see *Methods*) in order to investigate differential regulon activity (analogous to TF activity) within our dataset.

We first found that the IRF9 (interferon regulatory factor 9) regulon exhibited increased activity only in the Naïve IFN1 cluster (Fig. 2a), suggesting a role for the IRF9 TF downstream of type-1 interferon signaling and upstream of interferon-response gene transcription in B cells. In line with known literature, we also found that the XBP1^22^ and IRF4^22^ regulon activities were enriched in both ASC clusters (Fig. 2a). In murine models, XBP1 is a transcription factor necessary for terminal differentiation of plasma cells^23^ and, similarly, IRF4 is a transcription factor necessary for the generation of plasma cells^24^. Among other regulons, we also found enrichment of the ATF6 regulon in ASC clusters (Fig. 2a). ATF6 is involved in the unfolded-protein response and is activated by PRDM1^25^, another TF that controls plasma-cell development. Meanwhile, within the GC, GC IgA, and GC LMO2 clusters, we found upregulation of the regulon for the canonical GC transcriptional factor IRF8^8^ (Fig. 2a).

**Figure 2.**
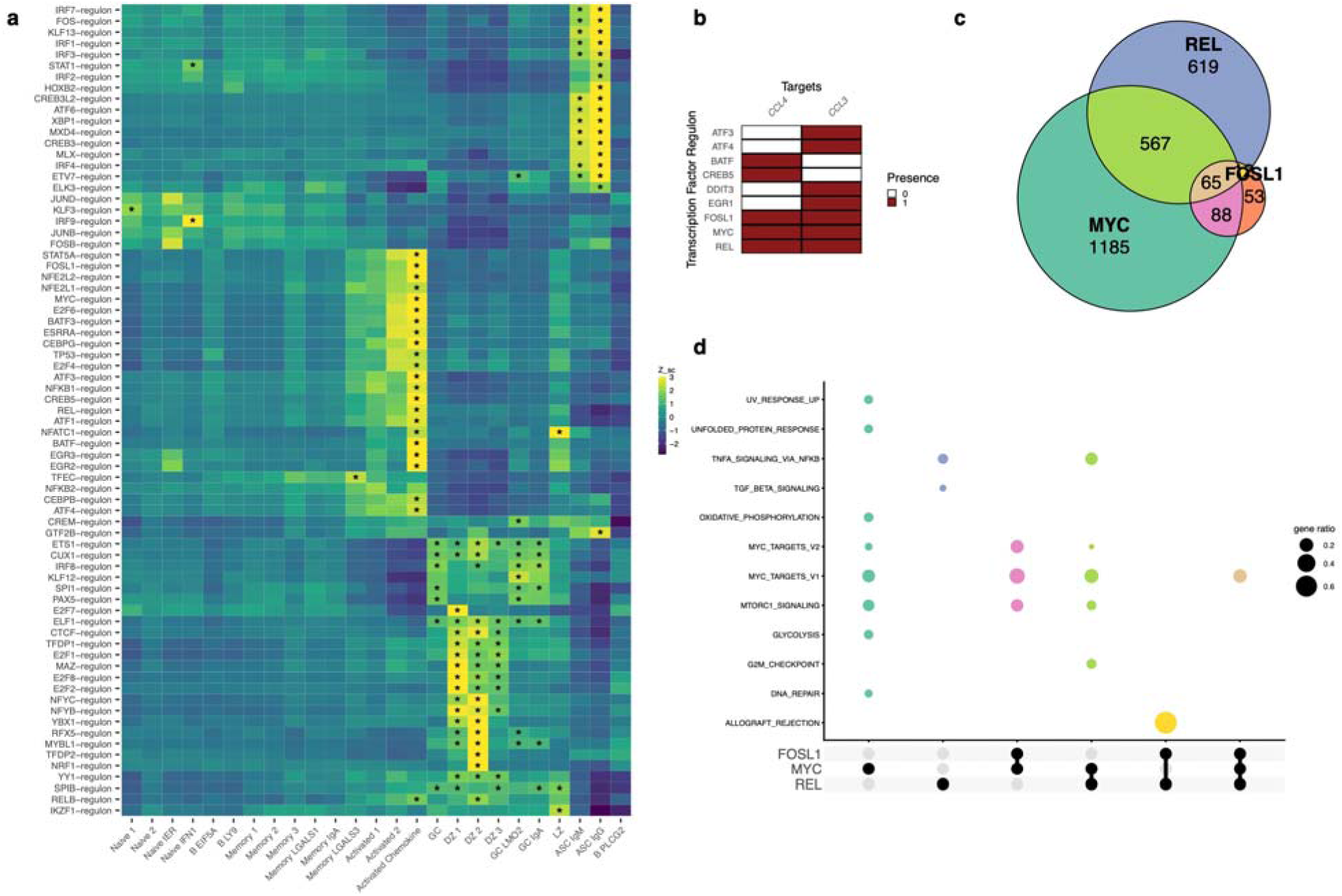
SCENIC analysis identifies MYC and REL as transcription factors predicted to regulate chemokine expression in activated B cells. **(a)** Average AUCell scores for selected regulons across all clusters. Regulons were selected for plotting if they were enriched (Wilcoxon rank sum test, AUC > 0.85) in at least one cluster. Heatmap cells were labelled with a star (*) if the geneset was enriched within that cluster. **(b)** Predicted targets for all regulons identified that target at least one of the *CCL4* or *CCL3* genes. **(c)** Venn diagram of the REL, MYC, and FOSL1 regulon targets. (**d**) Gene enrichment analysis of the various gene set intersections shown in panel **c** (hypergeometric test, *clusterProfiler*). Gene enrichment results only shown for enrichment tests with adjusted p-values < 0.05.

We next inspected the regulons enriched within the Activated Chemokine cluster, aiming to gain additional insight into the transcriptional processes driving chemokine expression. We found that this cluster showed overall enrichment for a relatively large number of regulons when compared to other clusters (Fig. 2a). We chose to identify the TFs whose regulons were predicted to target the chemokine genes *CCL4* and *CCL3*. Among chemokine-targeting TFs, we found that 3 TFs were predicted to target both of the chemokine transcripts: FOSL1, MYC, and REL (Fig. 2b); all three furthermore showed enrichment (AUC > 0.85) in the Activated Chemokine cluster (Fig. 2a, S4, **Table S5**). Among these regulons, both REL and FOSL1 shared considerable overlap of predicted downstream targets with MYC (Fig. 2c), suggesting overlapping targets for these three TFs.

We next partitioned the network structure of the REL, FOSL1, and MYC regulons to classify genes into 1 of 7 transcriptional programs (MYC-only, REL-only, FOSL1-only, MYC-REL, MYC-FOSL1, FOSL1-REL, FOSL1-MYC-REL, Fig. 2c). MYC-only targets were enriched for genes involved in metabolic processes (glycolysis and oxidative phosphorylation) along with cellular processes such as DNA repair and mTORc1 signaling (in addition to being enriched for genes from previously-derived MYC target gene sets) (Fig. 2d). On the other hand, REL-only genes had a target gene-set that was most enriched for genes involved in NF-kβ signaling and TGF-β signaling. (Fig. 2d). Enrichment analyses of the remaining gene sets (MYC-FOSL1, MYC-REL, FOSL1-REL, and FOSL1-MYC-REL) showed overlapping functions with the MYC-only and REL-only gene sets.

Overall, these enrichment analyses suggest that while the MYC and REL TFs both are predicted to target similar genes (including chemokine genes) and may have overlapping functions, there are particular processes, such as cellular metabolism, for which MYC may have a more overt regulatory role, and other processes, such as the downstream enactment of the NF-kB signaling pathway, in which REL may play a more overt regulatory role. In fact, the predicted regulon networks identify MYC as a target of REL and REL as a target of MYC (**Table S5**), underscoring a potential relationship between these two TF networks in BCR/CD40 activated human B cells.

### Gene expression and TF network pseudo-temporal modeling identifies genes and TF network expression changes associated with the GC to ASC transition

While the analyses above revealed identities and states of discrete B-cell populations, we considered whether we could explore the continuum of B-cell maturation using our scRNA-seq transcriptional level data. We chose to focus on GC to ASC maturation, and so we re-analyzed and re-clustered the GC and ASC clusters (Fig. 3a) independently of non-GC cells. We identified one GC cluster, GC LMO2 (which in fact was most similar to the same GC LMO2 cluster identified in the full analysis, Fig. 3b, Fig. S5), that showed a gradient of ASC marker expression (XBP1, PRDM1, MZB1, and IGHG4, Fig. 3b, Fig. S6) across its 2-dimensional UMAP representation, consistent with a cellular trajectory of differentiation from GC-like cells to ASC-like cells. We inferred that these cells captured a continuum of states across GC-to-ASC differentiation rather than a terminal cell type, and modeled transcriptional dynamics across this continuum based on these assumptions.

**Figure 3.**
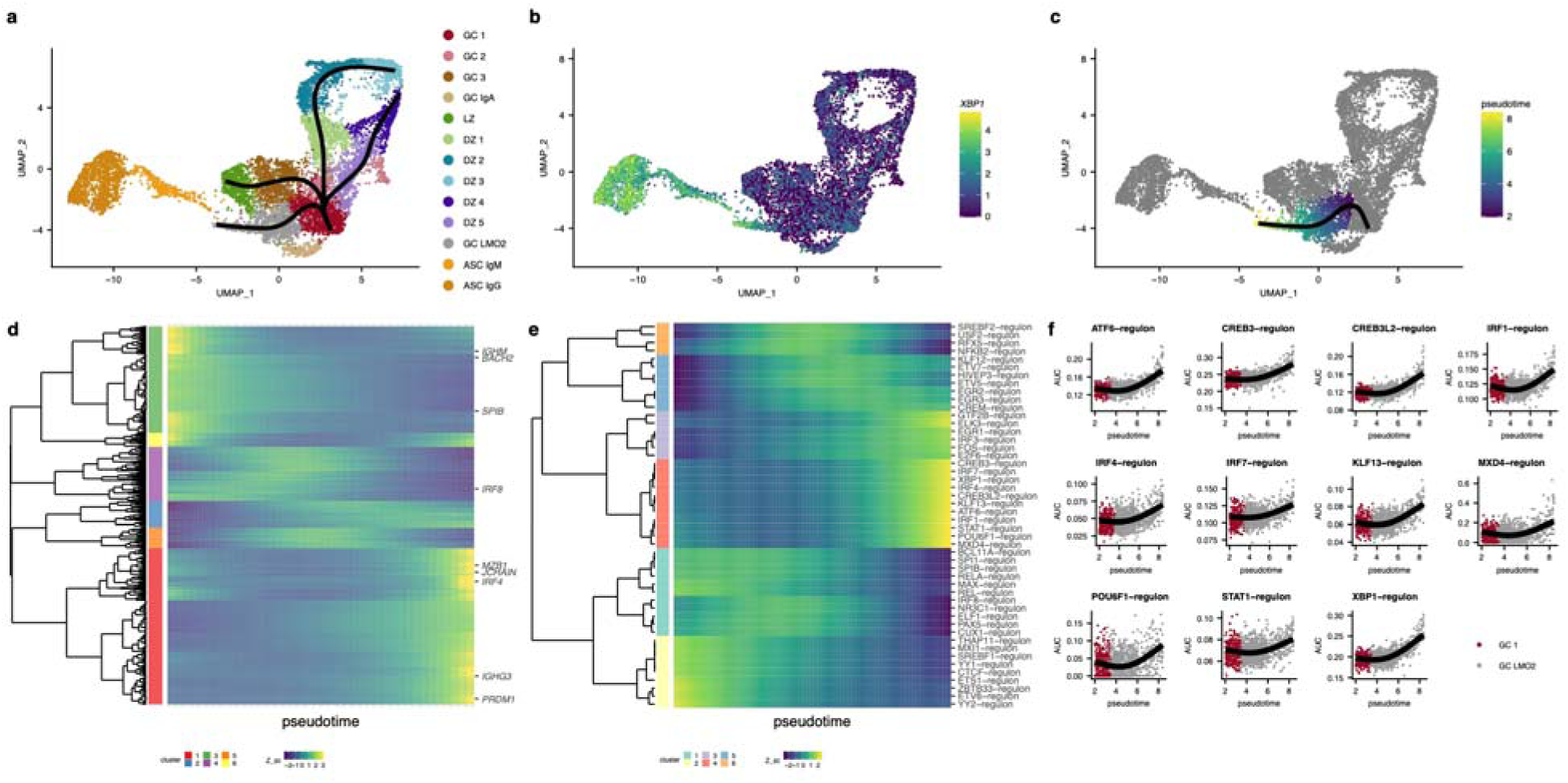
Trajectory inference of tonsillar B-cell scRNA-seq models transcriptional dynamics of a GC to ASC transition. (**a**) UMAP projection and cluster labels for the re-analysis (normalization, integration, dimensionality reduction) of the GC and ASC clusters. Slingshot trajectories (black) are overlaid on UMAP projection. (**b**) Gene expression levels of *XBP1* (log normalized counts) overlaid on cells in the UMAP projection. (**c**) Pseudotemporal ordering (pseudotime) results for trajectory of interest. (**d**) Modeled expression values across increasing pseudotime (left to right) for the cellular trajectory shown in panel **c**. Selected genes labelled. Expression patterns were clustered using Manhattan distance of the modeled gene expression values across pseudotime and hierarchical clustering was performed with a 6-cluster cutoff. (**e**) Modeled regulon activity values across increasing pseudotime (left to right) for the cellular trajectory shown in panel **c**. All regulons labelled. Expression patterns were clustered using Manhattan distance of the modeled gene expression values across pseudotime and hierarchical clustering was performed with a 6-cluster cutoff. (**f**) Generalized additive model results alongside regulon activity values are displayed for cluster 4 from panel **e**.

We leveraged the *slingshot*^26^ trajectory inference method and the *tradeSeq* gene expression modeling software to build *slingshot* differentiation trajectories on our UMAP projection (Fig. 3a) and order cells on the GC 1 to GC LMO2 trajectory (Fig. 3c) before modeling gene expression across this pseudotemporal axis. We then plotted the smoothed model expression values from the genes that with most significant gene expression variability over pseudotime (Fig. 3d, 967 genes with p-value < 0.01, full list of 10487 genes in **Table S6**) and identified 6 clusters of gene expression dynamics present in the data. One cluster (*red*, Fig. 3d) displayed an *increasing* gene expression pattern along the pseudotemporal axis. This cluster included canonical ASC genes such as IRF4, MZB1, PRDM1, JCHAIN, and IGHG3 (Fig. 3d). Two other clusters of genes (*green* and *purple*, Fig. 3d) displayed a *decreasing* gene expression pattern along the pseudotemporal axis, and included genes associated with the GC reaction such as *IGHM*, *IRF8*, *SPIB*, and *BACH2*^27^

In total, these gene expression patterns suggested that the pseudotemporal axis was consistent with a GC to ASC differentiation axis. We therefore chose to leverage our prior TF regulon approach to develop a method by which we could model TF dynamics in the GC-to-ASC transition. We considered that by modeling TF regulon-activity, rather than TF gene-expression, we may more optimally infer TF dynamics, assuming that predicted TF regulon activity (rather than TF gene expression) might be more consistent with true TF activity. Because TFs are known regulators of cell fate, this analysis may provide more salient targets for further study and/or inference. Using our TF pipeline in conjunction with our pseudotemporal modeling, we found a number of well-characterized GC-associated TFs, such as PAX5, IRF8, and SPIB^27^, that decreased in regulon activity along the GC to ASC transition (*green* and yellow, Fig. 3e). The canonical ASC TFs, XBP1 and IRF4^27^, instead increased in regulon activity along the transition (Fig. 3e-f). The ATF6, CREB3, and CREB3L2 TFs, which are associated with protein folding and cellular secretory function^28,29^, also increased in regulon activity along the GC to ASC transition (Fig. 3e-f). In addition to IRF4, two other members of the interferon-regulated factor family of TFs, IRF1 and IRF7, increased in regulon activity along the GC to ASC transition, and regulon activity of the STAT1 TF, a molecule activated upon interferon stimulation, also increased along this transition (Fig. 3e-f). Moreover, we identified a number of other TFs whose regulon activities increased along the pseudotemporal axis but whose roles have not been well described in GCs, ASCs, or B cells in general: KLF13, MXD4, and POU6F1 (Fig. 3f). Gene-level analyses of the TFs associated with increasing activity along the GC to ASC transition showed considerable variability and poor detection, in line with our postulate that TF regulatory network analyses would more optimally identify important TFs in cellular transitions (Fig S7).

## DISCUSSION

The antibody-independent functional capabilities of B cells in human health and disease remain incompletely defined. Here, we present a single-cell transcriptomic snapshot of human B-cell maturation and functional states within the human tonsil in order to infer potential B-cell functions beyond antibody secretion. Among our findings, we identify a dual CCL4 and CCL3 chemokine-expressing B-cell population with enrichment for a BCR/CD40 co-stimulatory transcriptomic signature, and implicate the transcription factors MYC, REL, and FOSL1 in regulation of this subset’s CCL4 and CCL3 chemokine expression. Furthermore, using a novel joint trajectory inference plus regulatory network computational approach, we model the transcriptional dynamics of the GC to ASC transition to identify associated candidate TFs involved in the GC to ASC transition. Overall, our data represents a resource for the probing of B-cell functional heterogeneity and maturation pathways within human lymphoid tissue and implicates associated transcription factors in driving the captured phenotypes.

The identification of a distinct B cell population expressing CCL4 and CCL3 concurrently with a transcriptomic profile of BCR and CD40 co-stimulation is consistent with prior reports demonstrating the potential for CCL4 and CCL3 production by in-vitro stimulated healthy naïve, memory, and GC B cells^30^ as well as malignant B cells^31,32^. While the ontogeny of these cells cannot be solely determined from our transcriptomic data, the lack of gene expression of typical GC markers (RGS13 and CD38, Fig 1d) in these cells and the distinct co-localization of the cells in the UMAP space with what are deemed non-GC cells (Fig 1b) suggest that these cells share more global transcriptomic similarities to cells outside of the GC reaction than to cells within the GC reaction. On the other hand, the enrichment of the LZ-gene signature (Fig 1e) within this population indicates these cells share some similarity to LZ cells. Further studies that recover B-cell repertoires in these cells and others with sufficient resolution within the secondary lymphoid tissues may provide insight into the ontogeny of these cells. Of further interest will also be determining where these B cells are detected spatially and temporally to GC structures (their presence in murine lymph node has been shown in ^4^) over the course of the immune response and their overall relation to the GC reaction. Our regulon analysis in particular highlights MYC as a potential TF driving expression of these chemokines, suggesting a temporal association of these B cells with pre-GC processes.

Outside of the CCL4 and CCL3 expressing B-cell population, we identify distinct expression patterns within memory B-cell subsets (LGALS1 and LGALS3) and activation patterns within B cells (type-1 interferon and early-activation gene expression patterns) that will serve to guide future studies into the role of these molecules in B cell function. Our regulon analysis further identifies candidate TFs implicated in the cellular identity of various clusters identified in our data that will serve as a useful complement to the growing high-throughput single-cell B-cell data in the literature^5,33,34^.

We present a novel approach for interrogating transcriptional dynamics along cellular differentiation pathway, which leverages an existing trajectory inference algorithm, *slingshot*^26^ but, rather than modeling gene-expression levels along the derived pseudotemporal axis, we instead leverage an existing gene regulatory network approach to infer TF-Target networks, and then model the activities of these networks along the pseudotemporal axis. We envision such an approach as being more capable of implicating TFs in cellular differentiation pathways, as the approach leverages TF-Target information, rather than TF gene expression, which may not adequately reflect TF activity.

A potential limitation of this approach is that certain TFs may share similar binding motifs which could implicate multiple candidate TFs when in fact, *in vivo*, only a subset of the identified TFs actually modulate a given expression program. We also note that our pseudotemporal approach relies on a-priori knowledge and assumptions for choosing the direction of the pseudotemporal axis. Nevertheless, we envision this approach as being a worthwhile addition to the growing tool-set used in single-cell transcriptomics and a useful tool for identifying candidate targets for further study.

Ultimately, we believe that the B cell transcriptional landscape analysis provided here will contribute to building a better understanding of B-cell function and maturation. We in fact transcriptomally characterize a chemokine-expressing B-cell population that potentially represents a pre-GC population of cells. In addition, our pseudotemporal modeling findings, which coincide with well-studied transcriptome dynamics involved the GC to ASC transition, further suggest a potential role for other, under-studied TFs. Furthermore, our regulon pseudotemporal modeling approach represents a viable alternative to gene expression based pseudotemporal modeling approaches, as incorporation of regulatory network provides stronger evidence for candidate TFs involved in cellular state transitions. We hope these analyses will provide a resource motivating further studies into the functional roles of specific B-cell subsets, as well as the roles and utility of specific molecules implicated in B-cell identity and maturation.

## METHODS

### 10X Chromium scRNA-seq

We isolated tonsillar mononuclear cells using ficcoll-graded centrifugation from 3 healthy human tonsil donors (4 year-old Male, 17 year-old Female, 5 year-old Male). From these, CD3-/CD14-/CD19+ live B cells were sort purified using FACS. Cells were rested overnight prior to single-cell RNA-seq using the 10X Chromium platform (v2 chemistry, 3’ end).

### scRNA-seq preprocessing and clustering

For bash, R, and python code implementation, see **DATA AVAILABILITY AND REPRODUCIBILITY.** Briefly, for our pipeline, we mapped scRNA-seq data to the human genome using CellRanger version 3.1.0. We next kept cells with at least 200 unique genes expressed and kept genes expressed in at least 3 cells. We then discarded cells with >5% read fraction deriving from mitochondrial counts, >4% read fraction deriving from a dissociation signature^35^, or >40,000 unique RNA features (Fig. S1A). We furthermore excluded contaminating non-B cells (predicted to be T cells based on detection of 2 or more of the markers CD3D, IL32 and CD2. We next used SCTransform^36^ to normalize and scale our data by regularized negative binomial regression (using 3000 variable genes per donor). We used the Seurat integration pipeline to integrate our data across the three donors using 40 CCA dimensions, followed by PCA on the scaled/integrated data, excluding immunoglobulin kappa/lambda chain genes (gene names beginning with IGKC or IGLC). We chose the first 35 PCs to build a kNN (30 neighbors, euclidean distance) from which we built a UMAP 2-dimensional projection and identified clusters using the Louvain algorithm (resolution = 1.3) on a shared nearest-neighbors graph derived from the kNN (minimum Jaccard distance prior to trimming = 1/15).

### scRNA-seq marker gene identification

We performed marker-gene identification using a one-vs-all approach for each cluster and a likelihood ratio test in which ‘donor’ was used as a latent variable in Seurat; those genes that showed an adjusted p-value < 0.01 and an average log2 fold change > 0.3 were considered ‘marker’ genes for each cluster.

### scRNA-seq geneset sources and module enrichment

We obtained light-zone and dark-zone signature genes from previous microarray profiling work on human tonsillar germinal center sorted B cells^37^. The G2M and S genesets were obtained directly from Seurat (‘cc.genes.updated.2019’). The MYC signature and the remaining Hallmark genesest were obtained from the Molecular Signature Database. Gene sets were scored at a single cell level using *AUCell,* part of the SCENIC toolset^21^.

### In-vitro peripheral B cell stimulation and bulk RNA-seq

Ficolled and FACS-sorted (CD3-/CD14-/CD19+) human peripheral B cells were stimulated with either BCR cross-linking antibodies (αBCR; Jackson Immunoresearch, Cata No. 109-006-129, 10ug/mL) or CD40L (Enzo Life Sciences, Cata No. ALX-522-110-C010, 1ug/mL) or the combination of αBCR and CD40L. After 18hrs, RNA was isolated from the differentially stimulated B cells using the RNeasy Plus Micro Kit (Qiagen) according to the manufacturer’s protocol. RNA quantity and quality were assessed by TapeStation and Nanodrop. Samples were run on a Bioanalyzer (Agilent) with the RNA 6000 Pico kit (Agilent) in order to calculate RNA integrity. Bulk RNA-seq was performed using a Novaseq6000 instrument (60 million reads/sample, 2×100bp, illumina). Reads were pseudoaligned to GENCODE reference genome v33 using kallisto^38^. DESeq2^39^ was used for differential gene expression analysis and a false-discovery rate of 0.05 was utilized. See **DATA AVILABILITY AND REPRODUCIBILITY** section for further kallisto and DESeq2 parameters utilized.

### Transcription factor gene-regulatory network analysis

We used the python implementation of SCENIC (*pySCENIC)* ^21^ on our full scRNA-seq count matrix. In brief, TF gene target networks are determined by construction of GRNs (using GRNBoost2, a random forest regression method) for each TF and pruning of their targets based on transcription factor motif enrichment. Activity of the resulting TF regulatory target networks, or regulons for short, was inferred on the whole single-cell dataset by using the AUCell algorithm, which scores the TF regulons independently for each cell, based on the enrichment for each regulon’s targets at the top of the gene expression rankings for each cell. Regulon activities were tested for differential enrichment in clusters using a Wilcoxon-rank sum test using the *Presto*^40^ package.

### Trajectory inference analysis

Transcriptomes belonging to the *GC* or *ASC* cluster partitions (Fig. 1) were re-analyzed from the *sctransform* step through marker-gene identification. For specific analysis parameters, please refer to **DATA AVAILABILITY AND REPRODUCIBILITY**. The resulting UMAP 2-dimensional reduction was used to construct differentiation trajectories using a modified form of *slingshot* as follows: (1) a pre-defined tree was defined on *GC* clusters such that a branch existed from the GC to GC-LMO2 cluster, (2) principal curves were built on this this branch using *slingshot*, (3) pseudotemporal ordering was performed setting *GC 1* as the root cluster. Gene expression-centric pseudotemporal analysis was performed using the *tradeSeq* package (knots = 3, see **DATA AVAILABILITY AND REPRODUCIBILITY** for full code), while regulon activity-centric pseudotemporal analysis was performed using the *mgcv* package (knots = 3, see **DATA AVAILABILITY AND REPRODUCIBILITY** for full code).

## DATA AVAILABILITY AND REPRODUCIBILITY

Complete R, python, and bash code (including R markdown notebooks with clear documentation) used to analyze and visualize our results have been deposited at https://github.com/diegoalexespi/espinozada-tonsil-paper-2021. Raw FASTQ files and processed CellRanger files for scRNA-seq have been deposited at GSE182221. Raw FASTQ files and processed kallisto files for bulk RNA-seq have been deposited at GSE182266.

## ACKNOWLEDGMENTS

DAE was supported by NIH Medical Scientist Training Program T32 GM07170 and T32 G000046. CLC and NR were supported by NIH NAID AI146026.

## AUTHOR CONTRIBUTIONS

DAE performed analysis and wrote the manuscript. CLC and RL performed experiments. NR, RL, and ABO supervised the project and edited the manuscript.

## FIGURES AND FIGURE LEGENDS

**Figure S1.**
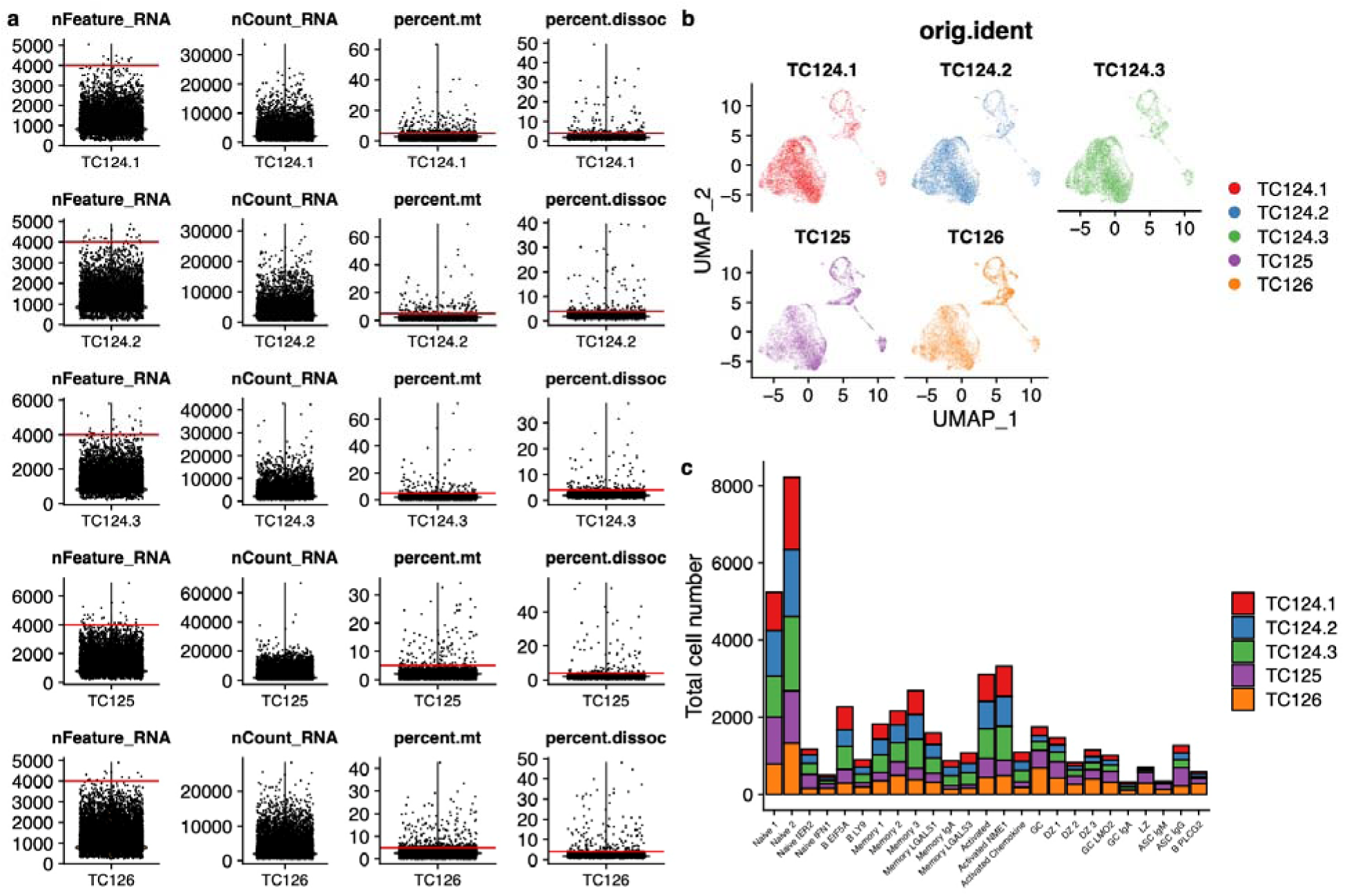
Preprocessing of scRNA-seq data. (**a**) Quality control metrics from five samples (3 biological replicates of TC124) with red horizontal lines denoting cutoffs used for downstream processing of scRNA-seq data. (**b**) UMAP projection of 45,376 single B-cell transcriptomes, colored and split by sample and/or donor. (**c**) Distribution of sample origin for cells within each cluster identified from Fig. 1c.

**Figure S2.**
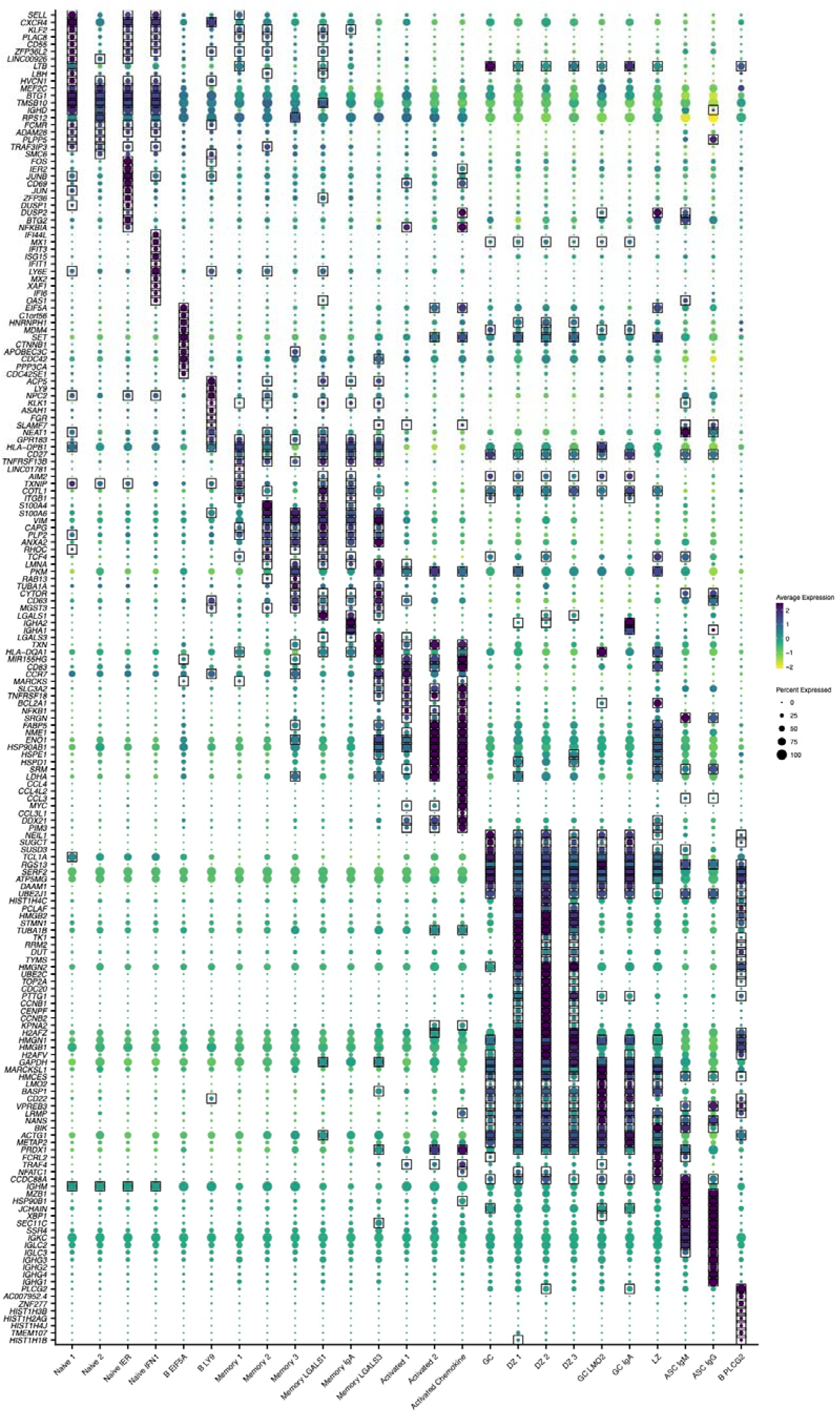
Average expression of top 10 marker-genes for each cluster. Top 10 marker-gene average expression (scaled log normalized counts) for each cluster. At least 10 marker-genes will be shown for each cluster. Genes were denoted as marker-genes if their average log2 fold-change was > 0.3 for the cluster of interest and adjusted p-value was < 0.01 (likelihood-ratio test). Marker-gene expression is denoted by square boxes on gene-cluster pairs.

**Figure S3.**
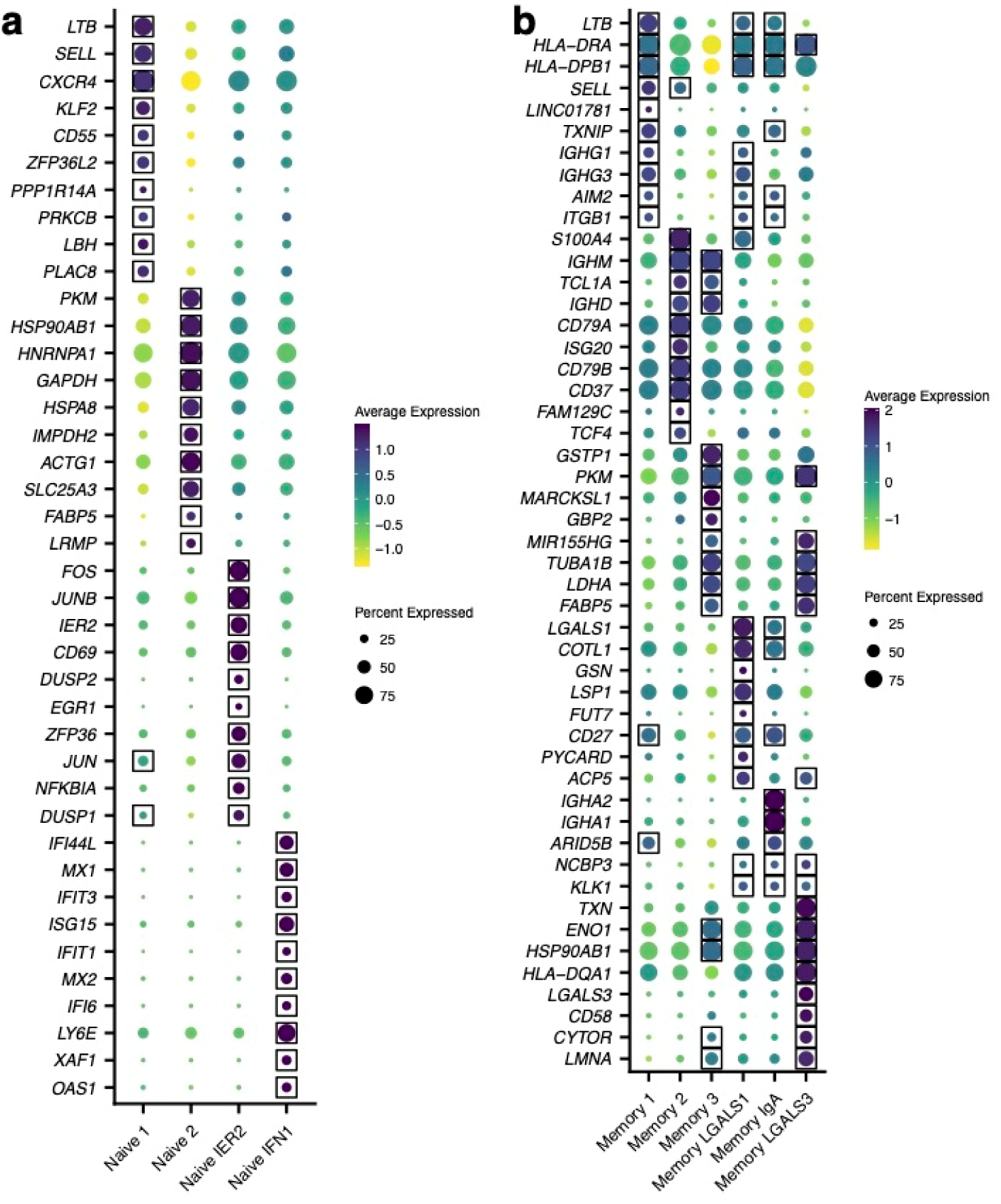
Memory-cell differential expression analysis. Marker-genes were determined for solely the memory-cell clusters (restricting comparisons to within the 6 clusters). Top 10 marker-gene average expression (scaled log normalized counts) for each cluster. At least 10 marker-genes will be shown for each cluster. Genes were denoted as marker-genes if their average log2 fold-change was > 0.3 for the cluster of interest and adjusted p-value was < 0.01 (likelihood-ratio test). Marker-gene expression is denoted by square boxes on gene-cluster pairs.

**Figure S4.**
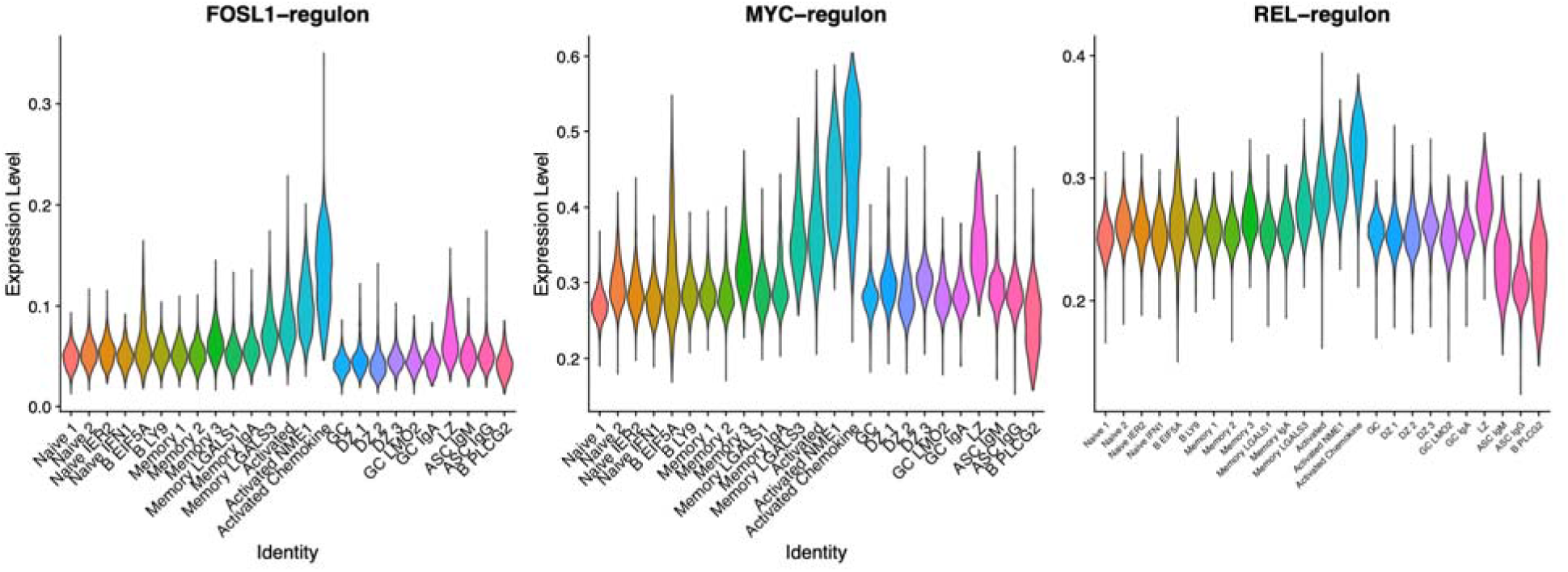
MYC, REL, and FOSL1 regulon activities across clusters. Violin plots of AUCell scores for the MYC, REL, and FOSL1 regulons across all clusters identified in Fig. 1b.

**Figure S5.**
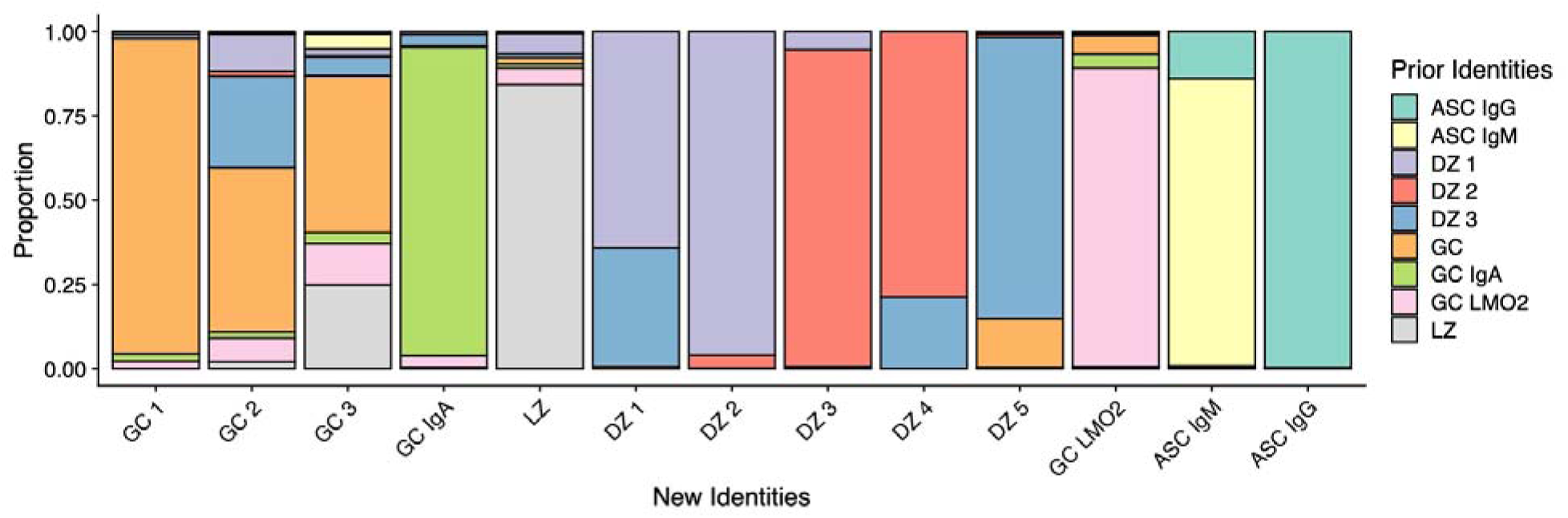
Proportions of original clusters in new clusters. Proportion of original cluster identities (from Fig. 1b) across new cluster identities (from Fig. 3a).

**Figure S6.**
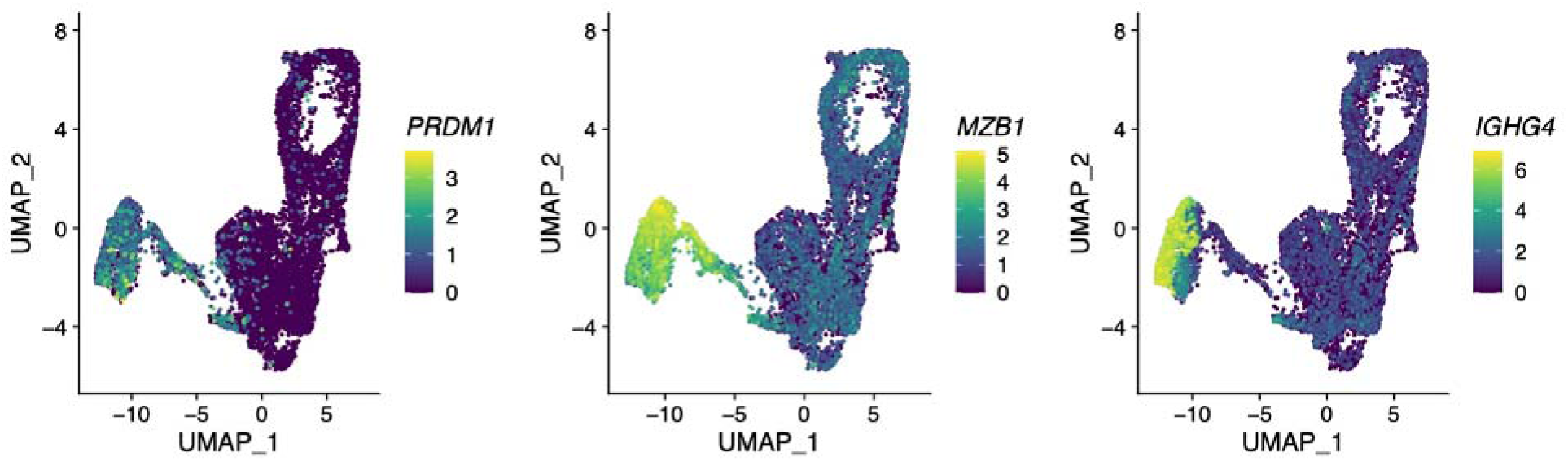
Gene expression levels of *PRDM1*, *MZB1*, and *IGHG4* in GC and ASC clusters. Gene expression levels of *PRDM1*, *MZB1*, and *IGHG4* (log normalized counts) gene expression overlaid on cells in the UMAP projection in Fig. 3a-b.

**Figure S7.**
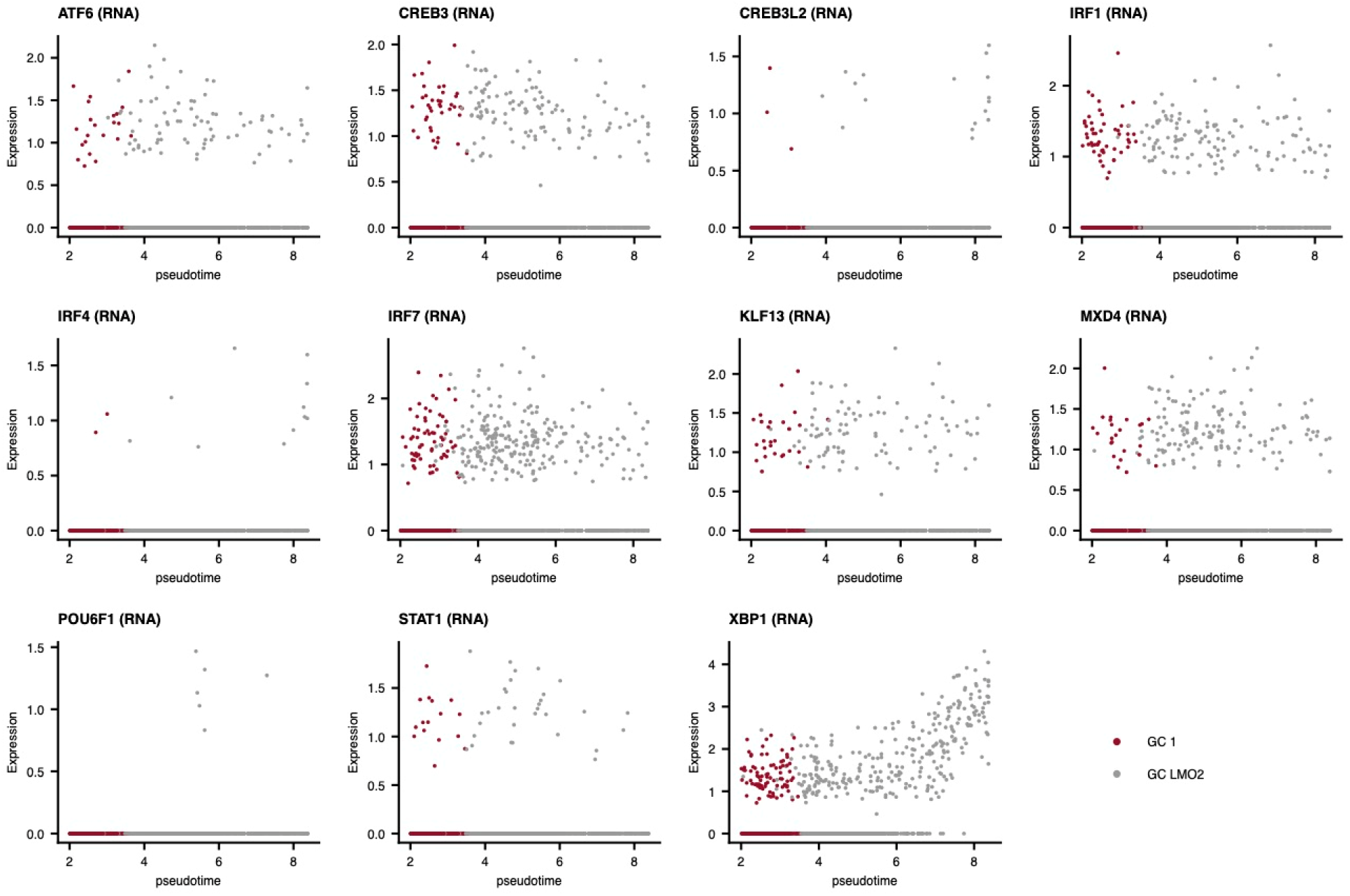
Gene-level expression of select TFs along the GC-to-ASC pseudotemporal axis. Gene expression levels along the GC-to-ASC pseudotemporal axis for the TF regulon activities plotted in Fig. 3f.

## TABLE LEGENDS

**Table S1. Donor information (n = 3) and cells recovered per 10X sample.**

**Table S2. Likelihood-ratio test gene differential expression results.**

**Table S3. Donor information (n=3) for each bulk RNA-seq sample.**

**Table S4. Identified gene signatures (BCR-only, CD40-only, BCR/CD40) from bulk RNA-seq.**

**Table S5. Wilcoxon rank sum test regulon differential expression results.**

**Table S6. *tradeSeq* association test results for GC to ASC trajectory gene differential expression.**

